# Psychometric and subcortical neurometric measures of temporal discrimination in rhesus macaques

**DOI:** 10.1101/2022.08.05.502987

**Authors:** Chase A. Mackey, Samantha Hauser, Adriana M. Schoenhaut, Namrata Temghare, Ramnarayan Ramachandran

## Abstract

Temporal envelope fluctuations are abundant in nature and are critical for perception of complex sounds. While psychophysical sinusoidal amplitude modulation (SAM) processing studies have characterized the perception of SAM, and neurophysiological studies report a subcortical transformation from temporal to rate-based code, no studies have characterized this transformation in unanesthetized animals or in nonhuman primates. To address this, we recorded single-unit responses and compared derived neurometric measures in the cochlear nucleus (CN) and inferior colliculus (IC) to psychometric measures of modulation frequency (MF) discrimination in macaques. IC and CN neurons often exhibited tuned responses to SAM in their rate and spike-timing. Neurometric thresholds spanned a large range (2-200 Hz Δ MF). The lowest 40% of IC thresholds were less than or equal to psychometric thresholds, regardless of which code was used, while CN thresholds were greater than psychometric thresholds. Discrimination at 10-20 Hz could be explained by indiscriminately pooling 30 units in either structure, while discrimination at higher MFs was best explained by more selective pooling. This suggests that pooled brainstem activity was sufficient for AM discrimination. Psychometric and neurometric thresholds decreased as a function of stimulus duration, but IC and CN thresholds were greater and more variable than behavior at durations less than 500 ms. This slower subcortical temporal integration compared to behavior was consistent with a drift diffusion model which reproduced individual differences in performance and can constrain future neurophysiological studies of temporal integration. These measures provide an account of AM perception at the neurophysiological, computational, and behavioral levels.

**Significance statement:** Listening in everyday environments tasks the brain with extracting information from sound envelopes. This process involves both sensory encoding and decision-making. Different neural codes for envelope representation have been well characterized in the auditory midbrain and cortex, but studies of the brainstem have usually been conducted in anesthetized rodents or cats. Moreover, these candidate neural codes have been studied in isolation from the decision-making process. In this study, we found that population activity in the primate subcortical auditory system contains sufficient information for discriminating sound envelope and applied a biologically plausible model of decision-making to sound envelope discrimination performance from rhesus macaques, a species with great phylogenetic and perceptual similarity to humans.

## INTRODUCTION

Temporal sound envelope fluctuations are critical for navigating complex acoustic environments (Bregman, 1994; Festen and Plomp, 1990; Vélez et al., 2012; Yost, 1991). Specifically, amplitude-modulation (AM) aids in the processing of species-specific communication sounds, including speech (Boemio et al., 2005; Drullman et al., 1994; Hauser, 1989; McDermott and Hauser, 2007; Richards and Wiley, 1980; Rosen, 1992; Zion Golumbic et al., 2013). Consequently, many studies have investigated AM processing at the psychophysical and neurophysiological levels. AM processing tasks have described human and nonhuman animal capacity for detecting and discriminating features of AM (Beitel et al., 2020; Dooling and Searcy, 1981; Fay, 1982; Formby and Muir, 1988; Kelly et al., 2006; Lee, 1994; Lemus et al., 2009a; Moody, 1994; O’Connor et al., 2011; Viemeister, 1979; Wakefield and Viemeister, 1990; Yao and Sanes, 2021). Primates and songbirds, specifically budgerigars, display enhanced perceptual sensitivity to AM relative to rodents (Kelly et al., 2006; Moody, 1994; O’Connor et al., 2011), positioning them as ideal models to investigate the neural basis of AM perception.

Neuronal responses to AM have been well characterized along the auditory pathway (Bartlett and Wang, 2007; Beitel et al., 2003, 2020; Bendor and Wang, 2008; Downer et al., 2017; Joris et al., 2004; Langner and Schreiner, 1988; Nelson and Carney, 2007; Niwa et al., 2013; Rhode et al., 2010; Sayles et al., 2013; Wang et al., 2008). In early stages (e.g. auditory nerve and cochlear nucleus) neurons respond with synchronized firing to AM, and in later stages (inferior colliculus and above) average rate-tuning becomes increasingly prevalent (reviewed in Joris et al., 2004). Consistent with this, the average neuronal sensitivity based on spike-timing in the cochlear nucleus (CN) and inferior colliculus (IC) correlated with AM detection performance (Henry et al., 2016; Sayles et al., 2013), while the average rate of simulated populations in auditory cortex (A1) correlated with AM detection performance (Johnson et al., 2012). Such population analyses have revealed that the decrease in spike-timing precision in auditory cortex (A1) may be recovered at the population level (Downer et al., 2021; Johnson et al., 2012), and can test predictions of modulation filter-bank models (Dau et al., 1997; Maxwell et al., 2020; Verhulst et al., 2018; Viswanathan et al., 2022), which factor heavily in current thinking of auditory perception.

Current knowledge of subcortical AM processing draws heavily from rodent, cat, and budgerigar studies; however, a potential concern is the lack of subcortical AM processing studies in primates. Nonhuman primates bear exceptional similarity to humans in CN and cortical neuroanatomical structure (Hackett, 2011; Moore, 1980, 2000; Moore et al., 1996; but see Rubio et al., 2008), perceptual measures of temporal processing (Kelly et al., 2006; Mackey et al., 2021a; Moody, 1994), auditory nerve frequency tuning (Joris et al., 2011; Verschooten et al., 2018), and cortical AM encoding (Hoglen et al., 2018). It is unclear to what degree the enhanced (relative to rodents) AM encoding in auditory cortex (Hoglen et al., 2018) is inherited from subcortical stations, because studies of subcortical AM processing in primates have been conducted under anesthesia (e.g. Rhode et al., 2010). Subcortical data from awake primates could enhance the understanding of human and nonhuman primate perception and speak to the validity of models suggesting that CN populations converge on IC neurons to confer AM sensitivity (Hewitt and Meddis, 1994; Nelson and Carney, 2004).

Finally, animal studies of the subcortical basis of AM processing have generally focused on detection paradigms, in contrast to cortical studies where the use of different paradigms has illuminated differences in their computational demands (Beitel et al., 2003; Lemus et al., 2009a, 2009b; Niwa et al., 2012; Yao and Sanes, 2021). These opportunities for furthering knowledge of the neural processing of AM were addressed in the present study through single-unit recordings in the CN and IC of awake macaques, and an AM frequency discrimination paradigm, in contrast to more typical detection paradigms.

## METHODS

### Subjects

Data from four adult male rhesus macaques (*Macaca mulatta*) are reported here. At the beginning of the study macaques ranged in age from 7-8 years old, with body weights ranging from 11-13 kg. Macaques in the behavior group were 7 (Monkey Is) and 8 (Monkey Da) years old at the beginning of data collection, and macaques in the electrophysiology group (Ch and De) were both 8 at the beginning of data collection. Macaques were fed a commercial diet (LabDiet Monkey Diet 5037 and 5050, Purina, St Louis, MO) supplemented with fresh produce and foraging items. Macaques were provided manipulanda as well as auditory, visual, and olfactory enrichment on a rotational basis. Macaques were fluid restricted for the study and received filtered municipal water averaging at least 20 ml/kg of body weight/day (typically closer to 25 ml/kg/day). Their weight was monitored at least weekly (typically 4 – 5 days each week) and stayed within bounds of the reference weights set by a veterinarian to index the animal’s health while on study. Macaques were maintained on a 12:12-h light:dark cycle and all procedures occurred between 8 AM and 6 PM during their light cycle. After repeated behavioral assessments to attempt to identify compatible social partners, these macaques were individually housed (due to incompatibility for social housing with available cohorts) with visual, auditory, and olfactory contact with conspecifics maintained within the housing room. The housing room was located in an AAALAC-accredited facility in accordance with the *Guide for the Care and Use of Laboratory Animals*, the Public Health Service Policy on Humane Care and Use of Laboratory Animals, and the Animal Welfare Act and Regulations. Macaques in this colony received routine health assessments including twice-yearly tuberculosis testing and updating of reference weights. All research procedures were part of protocols that were approved by the Animal Care and Use Committee at Vanderbilt University Medical Center (VUMC).

### Surgical procedures

Monkeys were prepared for chronic experiments using standard techniques employed in nonhuman primate studies and as reported in previous studies (Dylla et al., 2013; Rocchi and Ramachandran, 2018, 2020). Briefly, anesthesia was induced via administration of ketamine and midazolam, and maintained via isoflurane. A headpost (stainless steel/plastic) was implanted on the skull to restrict head movement during experiments and minimize sound pressure level variability at the ear as a result of positioning across behavioral sessions. The headpost was secured to bone using 8 mm titanium screws (Veterinary Orthopedic Implants, St. Augustine, FL) and encapsulated in bone cement (Zimmer Biomet, Warsaw, IN). Multimodal analgesics (pre- and post-procedure), intra-procedure fluids, and antibiotics (intra-procedure) were administered to the monkeys under veterinary oversight. The other two surgeries implanted recording chambers (Crist Instruments, Hagerstown, MD) on the skull around craniotomies at stereotaxically guided locations. The recording chambers were angled to fit on the skull – the midbrain chamber was tilted lateral 20° and the brainstem chamber was angled posterior 26°. The chambers were chosen with bases that fit the cranial curvature and were secured to the skull using bone cement and screws. Intra-procedure fluids were provided, and pre- and postsurgical analgesics were administered, and the monkey was monitored carefully until complete recovery had occurred.

### Apparatus and Stimuli

Monkeys were seated in a primate chair, designed and constructed in-house, and situated inside a sound treated booth (IAC, model 1200A). Stimuli were presented in the free field via a speaker (Rhyme Acoustics, NuScale 216, available through Madisound - Middleton, WI) located in the frontal field 90 cm from the center of the monkey’s head. Broadband noise was 76 dB SPL, 500 ms in duration (unless otherwise specified), ramped on/off using a 20 ms cosine squared function (10 ms for durations less than 250 ms), and sinusoidally amplitude-modulated at different frequencies (2-1024 Hz). The speaker was calibrated with a ¼ inch microphone (378C01, PCB Piezotronics) positioned at the location where the entry to the monkey’s ear canal would be during experiments. The speaker was calibrated to ensure that outputs were within 3 dB across all frequencies.

### Task design

Experimental flow was controlled by a computer running OpenEx software (System 3, TDT Inc., Alachua, FL). Monkeys Is and Da performed a lever-based reaction time Go/No-Go discrimination task (Figure 1A). Trials were initiated by pulling a lever, following which a delay of 1.5-2 seconds elapsed and two noise bursts modulated at 100% depth were presented without an interstimulus interval (ISI). Trials could be signal trials (50%) in which two AM noise bursts of differing AM frequency were presented, or they could be catch trials (50%), during which identical AM noises were presented. Lever release was required within a response period of 1 second (after the second noise onset) to indicate discrimination on signal trials, and animals were required to continue to hold the lever on catch trials. All reward contingencies followed convention established in previous studies (e.g. Beitel et al., 2003, 2020; Moody, 1994). Lever release on signal trials (hits) were rewarded with fluid. Lack of lever release within 1 second of the onset of the second noise onset on signal trials (misses) was punished with a time-out of 4-6 seconds in which no trial could be initiated. Lever release on catch trials (false alarm) also resulted in a 4-6 s timeout. Correct rejections (lack of release on catch trials) were rewarded with fluid. Rewarding correct rejections and punishing misses was necessary to balance reward contingencies across trial types, and we empirically determined these reward contingencies to be optimal for performance. The green and red in Figure 1A indicate fluid reward and time-out punishment, respectively. MF was varied trial-to-trial, over a range of 1-64 Hz Δ MF. Each MF was repeated 30 times, resulting in blocks containing ∼300-420 trials. Experiments were performed after confirming normal hearing status through extensive audiometric and physiological characterization (ABRs, DPOAEs, otoscopy, and tympanometry), ensuring that all measures were within the range of normative values as reported in our previous work (Burton et al., 2019; Hauser et al., 2018; Stahl et al., 2022; Valero et al., 2017). Figure 1B shows how monkeys Is and Da learned to discriminate amplitude modulation frequency (MF) at various MFs in about ten sessions. These data were collected using a cage-training apparatus. A speaker, lever, and waterspout were mounted to the front of the primate cage for short periods each day, permitting the monkeys to learn to interact with the lever for a food reward. Gradually, monkeys were transitioned into pulling the lever for fluid, and, eventually, to releasing the lever for signal trials, but not catch trials.

**FIGURE 1.**
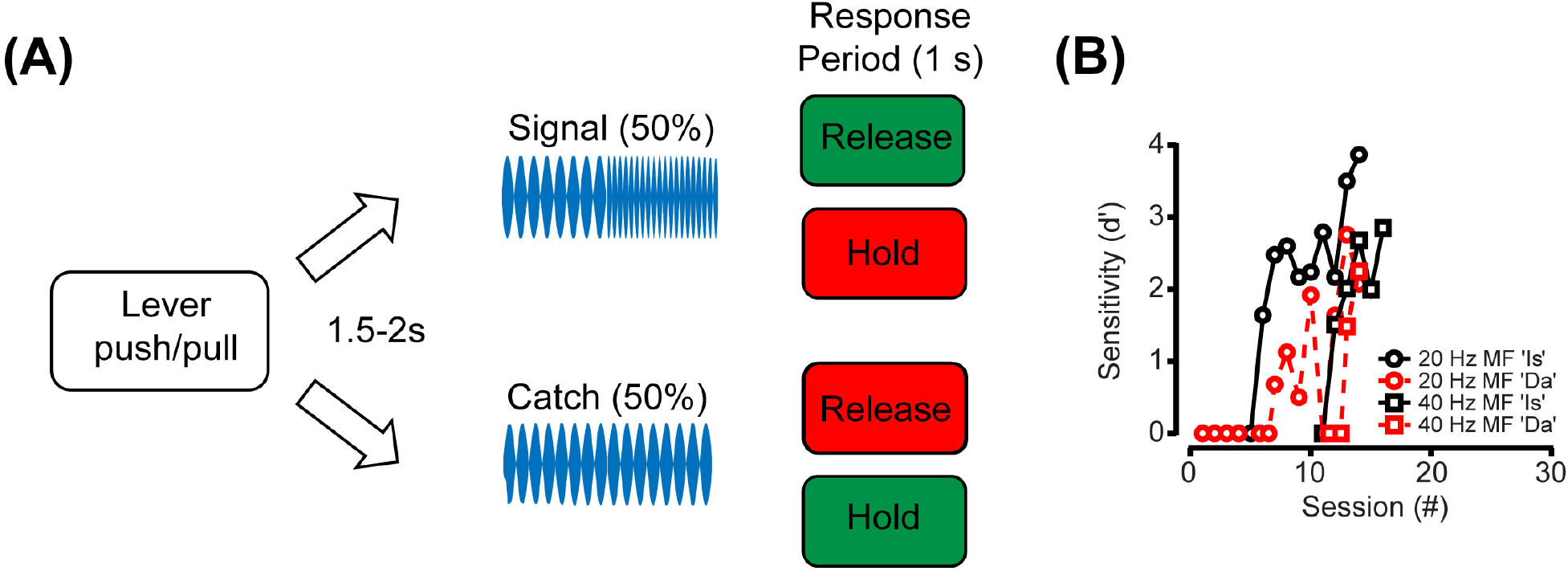
Task design and learning trends. **A**. A diagram of the go/no-go procedure used to measure temporal discrimination performance in this study (see Methods). Monkeys initiated a trial by pulling a lever. Following a random delay (1.5-2 seconds), stimuli were presented for a signal or catch trial (50/50). Monkeys were rewarded (green boxes) for lever release within the response period (1 second) of a signal trial (“Hit”) or holding on catch trials (“Correct rejection”). Monkeys were punished with a timeout (red boxes) on holding the lever through the response period of a signal trial (“Miss”), or for releasing on a catch trial (“False alarm”). **B**. A signal detection theoretic measure of sensitivity (*d’*) is shown as a function of training session for each monkey (shown in different colors). Over the course of training, discrimination performance increased for both monkeys discriminating either 20 Hz (circles) or 40 Hz amplitude-modulated noise (squares).

### Neuronal recordings

Monkeys Ch and De (the electrophysiology group) sat passively during recordings reported here, though they had experience in previously published tone detection tasks, including AM processing tasks (Bohlen et al., 2014; Mackey et al., 2021b). Single-unit recordings were made in the CN (n = 35) and in the tonotopic portion of the IC (n = 34). A glass coated tungsten electrode (Alpha Omega Engineering, Alpharetta, GA; tip length ∼7-10 μm, diameter ∼5 μm; Thomas Recording GmbH, tip impedance 2 – 4 MΩ) was placed in a stainless steel guide tube near the surface of the brain. The guide tube was advanced manually 10 mm into the brain when approaching the central nucleus of the inferior colliculus (IC; all reference to IC in text and figures refers to the central nucleus) and about 15 mm when approaching the cochlear nucleus (CN). The electrode was advanced further into the brain by means of a remotely controlled hydraulic micromanipulator (MO-97, Narishige Inc., Hampstead, NY). The electrode traveled through the cortex to reach the IC, and through the cortex and cerebellum to reach the CN. As the electrode was driven into the brain, bursts of noise were used as probe stimuli to assess proximity to auditory structures. Proximity to CN or IC was indicated by changes in background responses to the noise bursts. The IC was identified as the auditory structure posterior to the superior colliculus (identified by superficial visual drive, deep layer eye movement sensitivity) and anterior to the cerebellum (identified by simple and complex spikes). Further criteria to identify IC (after Nelson et al. 2009) included: (i) short latencies (≤ 20 ms); (ii) reliable, non-habituating responses; and (iii) identification of presence in tonotopic gradient, encountered by several groups, including the present authors (Nelson et al., 2009; Rocchi and Ramachandran, 2020; Ryan and Miller, 1978; Shaheen et al., 2021). The CN was identified as the region of auditory responses medial to the flocculus (identified by simple and complex spikes, and eye movement sensitivity to ipsiversive and downward eye movements observed over the video monitor). Histological verification of the locations of electrode penetrations in the IC and CN were reported previously (Mackey et al., 2022; Rocchi and Ramachandran, 2020).

Once the electrode moved into the CN or IC, single units were isolated using tones (Ramachandran et al., 2000; Rhode et al., 2010; Rocchi and Ramachandran, 2018). The electrode was advanced through the CN or IC until the signal from the electrode was predominantly from one unit. Single units were verified based on visual inspection, and principal component analysis available in the TDT System 3 software suite. The filtered waveform of the electrode signal and the waveforms of spikes that exceeded a user-defined threshold were sampled at 24.4 kHz and stored for offline analysis. Once a single unit was isolated, its characteristic frequency (CF: the frequency with the lowest threshold) was estimated and used to derive the frequency tuning of the unit via a frequency response map (FRM). A FRM was obtained by measuring the responses of the unit to tones as a function of frequency and sound pressure level. Frequency was varied over a 2 or 4 octave range around the estimated CF in 100 logarithmically spaced steps and at three or four sound levels, starting near CF threshold values, and proceeding in 20 dB steps. FRMs were also used to classify the response type of the CN or IC units (Evans and Nelson, 1973; Ramachandran et al., 1999). Sound pressure levels used for the FRMs ranged from 10-15 dB below estimated threshold to as high as 74 dB SPL.

### Calculation of responses to amplitude-modulation

Average firing rate was calculated over the entire 500 ms interval that each AM noise burst was presented, at each modulation frequency. Changes in firing rate relative to the response to steady-state noise were plotted, depicting the rate modulation transfer functions (MTFs) as others have previously reported (Krishna and Semple, 2000; Langner and Schreiner, 1988; Nelson and Carney, 2007; Sayles et al., 2013). Spike-timing was measured using classical vector strength (Goldberg and Brown, 1969) and phase-projected vector strength (Yin et al., 2011). The onset response was included for both firing rate and vector strength (VS) calculation (see Discussion).

### Calculation of accuracy and speed

Detection theoretic methods were used to analyze both behavioral and neuronal responses (Macmillan and Creelman, 2004; Swets, 1973; Tanner and Swets, 1954). Behavioral performance from each block of data was analyzed to calculate hit rate at each modulation frequency (*H(MF)*) and false alarm rate (*F*). Sensitivity at a given modulation frequency (MF) was calculated as *d*′
s(*MF*) = (z(*H*(*MF*)) − z(*F*)+, where *z* represents calculation of the z-score of the value, implemented in MATLAB via the function “norminv.” From *d’*, probability correct in a two-alternative forced-choice experiment was given by *pc(MF)*, as *pc*(*MF*) = z^−1^(*d*′(*MF*) / 2), where *z*^*-1*^ represents the transformation from a standard normal variate to probability correct. Calculation of *pc*, rather than a more common *d’* measure, permitted comparison with our distribution free (ROC) calculation of probability correct based on neuronal responses (Mackey et al., 2022; Rocchi and Ramachandran, 2018, 2020). Psychometric and neurometric functions were fit with a modified Weibull cumulative distribution function (CDF) as others have done in both detection and discrimination tasks (Britten et al., 1996; Christison-Lagay and Cohen, 2014; O’Connor et al., 1999). The modified equation was:

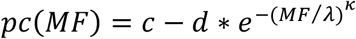

where *c* represents saturation and *d* represents the range of the function, and *λ* and *κ* represent the threshold and slope parameters respectively. The threshold was calculated as that tone level at which *pc*_*fit*_*(MF)*=0.76, similar to common use of *d’=1* threshold criterion. Reaction times (RT) are reported from hit trials and were calculated as follows:

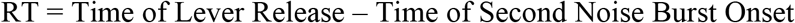

RTs were plotted cumulatively to visualize and to give insight into the effects of modulation frequency difference on discrimination speed, as previous publications of AM processing in animals have rarely reported RTs.

### Simulation of population responses

Neuronal responses were analyzed at the population level in a similar fashion to the “across-cell” method described by Johnson et al. (2012). A population (2-35) of neurons was randomly selected with replacement, following which single-trial spike trains were randomly sampled from each neuron. Responses were sampled 20 times with replacement, after determining empirically that neurometric sensitivity could not be enhanced beyond 20 permutations. Rate responses were summed, while vector strength was calculated based on the pooled spike times. The pooling process was repeated 1,000 times. The simulated population response to each modulation frequency calculated in this way served as the basis for ROC calculation described above, permitting the calculation of the population’s neurometric probability correct and discrimination threshold. Pooling is not reported for vector strength, as we found that VS only decreased as spike-trains were pooled, likely due to the diverse phase-preferences of subcortical neurons.

### Time-window analysis

Neuronal responses were calculated at varying durations to compare neuronal sensitivity to behavioral sensitivity as a function of stimulus duration. This was done by changing the time-window within which spikes were counted. This was initially attempted at exactly the durations used in the behavioral experiments (50, 100, 250, and 500 ms), however, the concentration of spikes at the onset of each cycle of AM in the noise caused highly variable and non-monotonic spike-counts. We then calculated time-windows differently for each modulation frequency, so that the time-window would be rounded up or down to the nearest AM trough. Using this method, and slightly different durations (50, 62.5, 125, 250, 500 ms), more consistent responses were attained as a function of duration. After the time-window was adjusted, ROC analysis was conducted on the spike-count using the method described above, and in our previous publications (Mackey et al., 2022; Rocchi and Ramachandran, 2018, 2020).

### Drift-diffusion model

Data were fit with a drift diffusion model using the HDDM 0.8.0 package (Wiecki et al., 2013), based on previous work modeling Go/No-Go task performance (de Gee et al., 2020). We used some of the code made available by de Gee et al. on Github (https://github.com/hddm-devs/hddm/blob/master/hddm/examples/gonogo_demo.ipynb). Data were fit using RT quantiles, using the G-square method. RTs, along with proportion of go responses at each modulation frequency and duration, contributed to G square, and a single bin of the number of no-go responses (but not the no-go RTs) also contributed to G square, which is conventional for Go/No-Go tasks (de Gee et al., 2020; Ratcliff et al., 2018). Drift rate was allowed to vary across modulation frequency, while non-decision time and decision-threshold could only vary as a function of whether the trial was a catch trial or not, consistent with the finding that high conflict trials can induce rapid changes in these processes (Cavanagh et al., 2014). This model was chosen based on a Bayesian information criterion (BIC) value that was lower than the BIC of other models that were less psychologically plausible (e.g. ones in which drift rate, threshold, and non-decision time could vary across modulation frequency, ones in which drift rate and threshold could vary across modulation frequency, and ones in which drift rate was the only free parameter). All data were fit using Markov-chain Monte Carlo sampling, using 6000 samples and a 1000-sample burn-in. The goodness of fit of the model was quantified with BIC (discussed above), and with a posterior predictive check, which indicated that the 95% confidence intervals around the model estimates of the data captured both the empirical choice proportions and empirical reaction-time data.

### Statistical analyses and curve-fitting

In all cases, curve fits were attained via non-linear least squares method implemented in MATLAB 2019a (Mathworks Inc.). Two-sample Kolmogorov-Smirnov tests were used to statistically confirm the differences in neurometric threshold distributions (“kstest2” in MATLAB). Statistical analysis of the goodness of fit of the drift diffusion model is discussed above.

## RESULTS

### Psychometric measures of temporal discrimination

Psychometric measures of discrimination were attained through a Go/No-Go procedure (see Methods, Figure 1A). After learning to discriminate between noises with large differences in modulation frequency in their home cages (see Methods, Figure 1B), macaques were trained to discriminate AM frequency near discrimination threshold in a sound booth. Figure 2 shows psychometric functions (Figure 2A-3D) and reaction-time distributions (Figure 2E, F) from Monkeys Is and Monkey Da. Hit rate and false alarm rate were used to calculate a signal detection theoretic measure, Probability Correct (PC, see Methods). PC was used in favor of more typical measures such as *d’* to facilitate comparison to neurometric measures, consistent with previous studies from this group (Burton et al., 2018; Hauser et al., 2018; Mackey et al., 2022; Rocchi and Ramachandran, 2018, 2020). The psychometric functions and RT distributions in Figure 2E and F extend what is currently known about MF discrimination in animals, as previous studies have usually reported only thresholds attained through adaptive procedures. Figure 2 shows, for the first time, that response accuracy and speed change coincidently in this task, indicating that performance improved as the change in MF increased.

**FIGURE 2.**
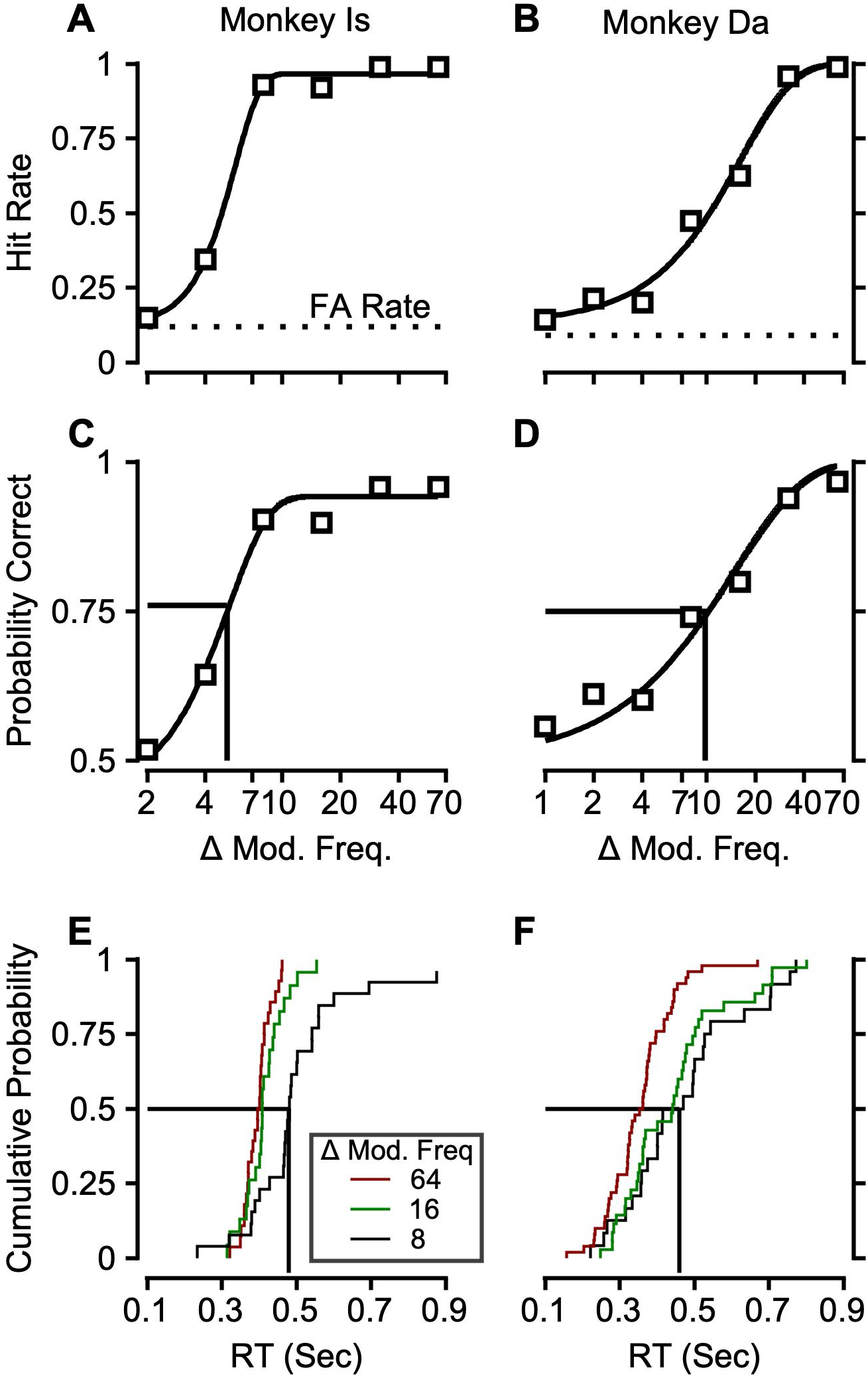
Amplitude modulation frequency discrimination performance from the two monkeys. **A**. Hit rate and false alarm rate for monkey Is, plotted as a function of change in modulation frequency (MF) of a 20 Hz, broadband, amplitude modulated noise. **B**. Hit rate and false alarm rate for monkey Da. **C, D**. A signal detection theoretic measure of sensitivity (Probability Correct, see Methods) calculated from the hit rate and false alarm rate in **A** and B, plotted as a function of change in MF, for monkeys Is (**C**) and Da (**D**). Psychometric functions were fit with a Weibull cumulative distribution function (see Methods), and a conventional value of PC = 0.76 was used as threshold criterion. **E, F**. Reaction times plotted cumulatively at example modulation frequencies for monkeys Is (**E**) and Da (**F**).

The psychometric thresholds attained from monkeys Is and Da are nearly identical to previous reports in macaques (Moody, 1994). Moody (1994) also synthesized and reported data from other species. Figure 3 shows discrimination thresholds from those previous studies (black) compared with data in the present study (color).

**Figure 3.**
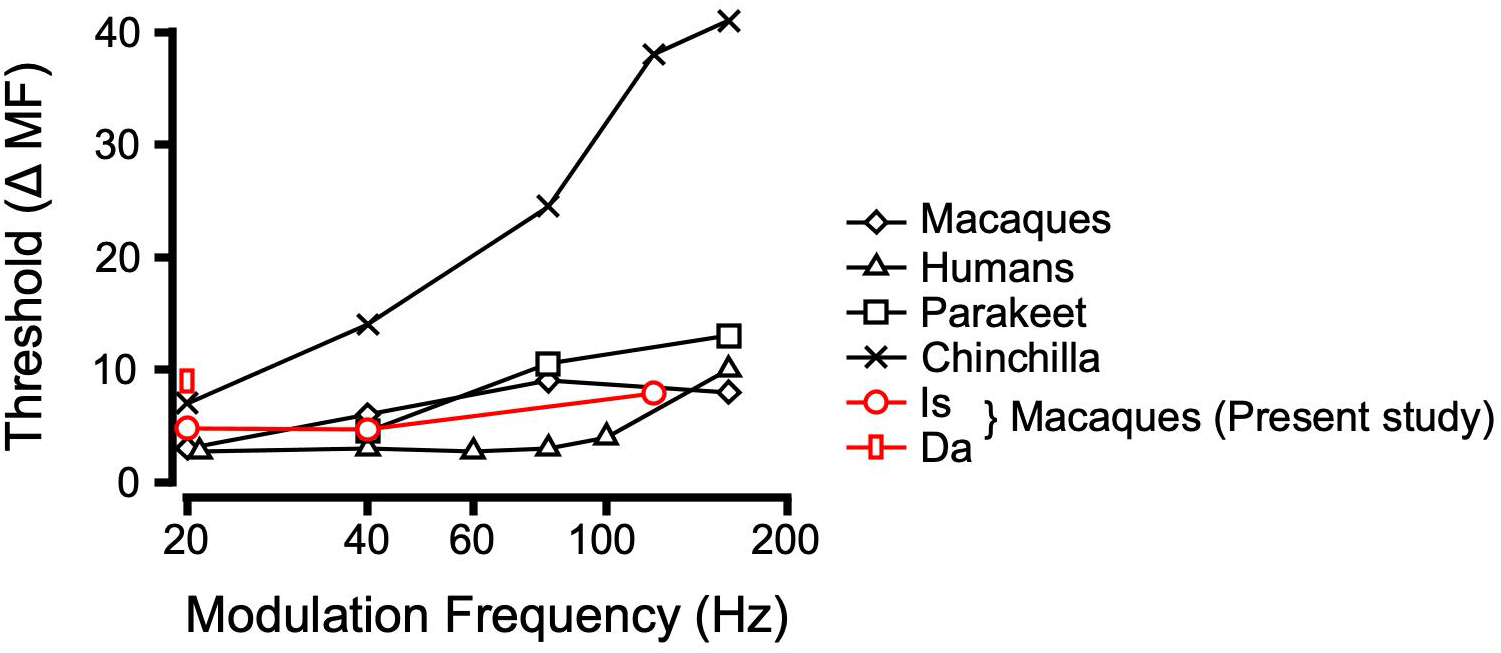
Amplitude modulation frequency discrimination thresholds as a function of the standard frequency across species. Macaque discrimination thresholds from the present study are plotted along with data from a variety of species originally reported by Moody (1994). Monkey Is completed psychometric functions (from which thresholds were extracted) at 20, 40, and 120 Hz, while monkey Da only completed psychometric functions in the 20 Hz condition. Data from the present study are shown in red (circles for Monkey Is, rectangles for Monkey Da).

### Modulation Transfer Functions in the Cochlear Nucleus (CN)

Initial characterization of how single neurons in the auditory pathway respond to temporal sound envelope fluctuations often takes the form of modulation transfer functions (MTFs). MTFs are displayed as changes in average firing rate, or spike-timing (measured using vector strength), as a function of modulation frequency (see Methods). The present report documents the first MTFs in the subcortical auditory system of awake primates, from Monkeys Ch and De, (see Methods). Figure 4 shows example MTFs from the cochlear nucleus (CN). Figure 4 shows units that exhibited typical changes in vector strength as a function of MF, while Figure 5 shows units that exhibited changes in firing rate as a function of MF, which were encountered more frequently than expected. A limitation of this study is that only 35 neurons were recorded in the CN, which is a necessary caveat in interpreting the proportions of each form of MTF in the sample. However, the entire tonotopic axis was sampled, shown by the distribution of characteristic frequencies as an inset in the top panel of Figure 4. This sample contained mostly band-pass and band-reject temporal MTFs. Figure 4 contains the specific percentages of each temporal MTF form. Interestingly, we found many examples of rate-tuning to AM in the putative VCN (based on frequency response maps being exclusively Type I and III in our sample). Figure 5 shows examples of these rate MTFs.

**FIGURE 4.**
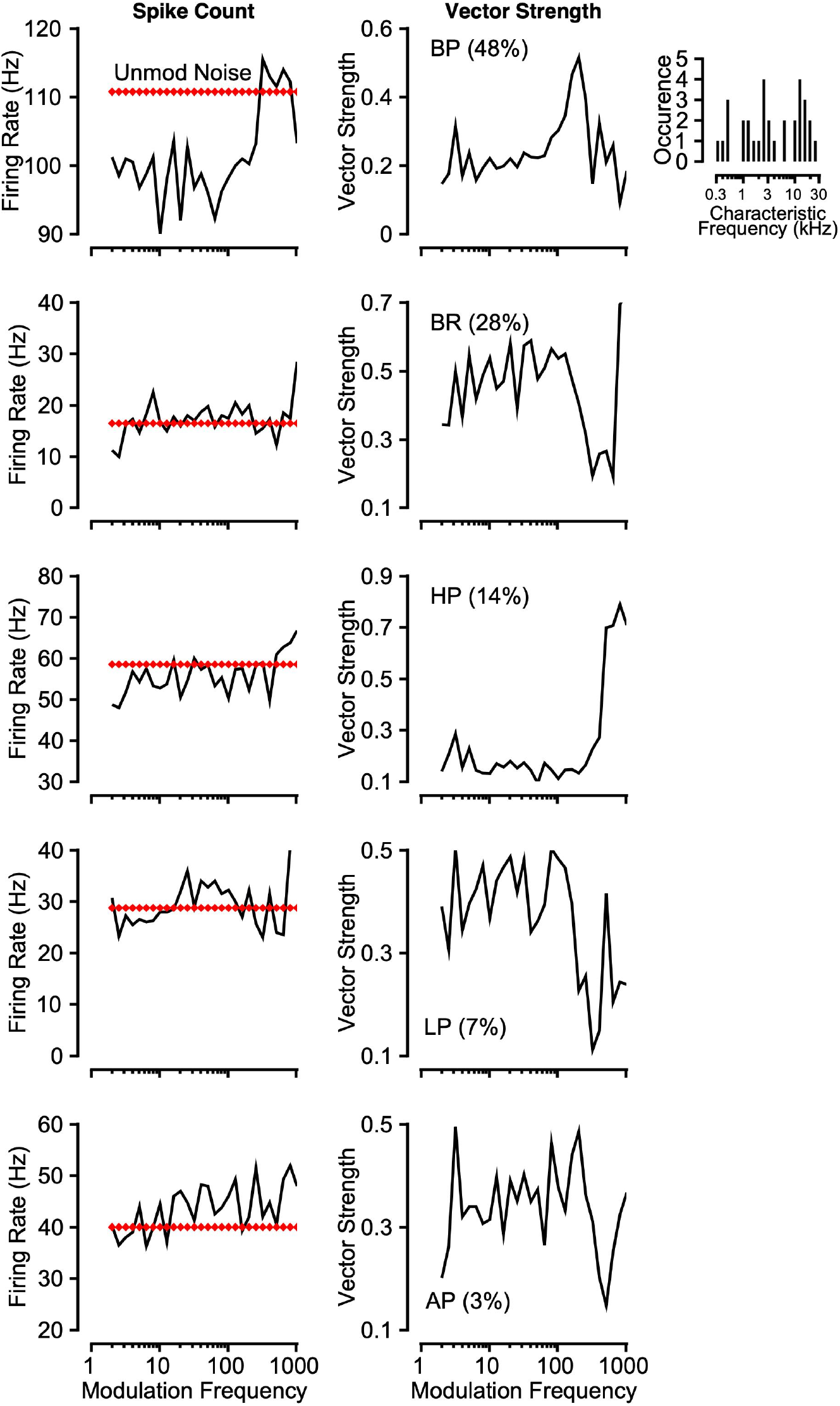
Modulation transfer functions (MTF) in the cochlear nucleus. Each row shows an example neuron’s responses to amplitude-modulated noise in terms of rate (left) and temporal metrics (right) to represent rate and vector strength (VS) based MTF (black lines). Dotted red lines shows the rate response to unmodulated noise. A histogram of the characteristic frequencies encountered is shown as an inset as evidence that most of the tonotopic axis was sampled. Neurons were categorized based on their VS-MTF shape, and exhibited conventional shapes (Band-pass, band-reject, high-pass, low-pass, all-pass). The percentage of each VS-MTF shape encountered is displayed in the figure legend of each panel.

**FIGURE 5.**
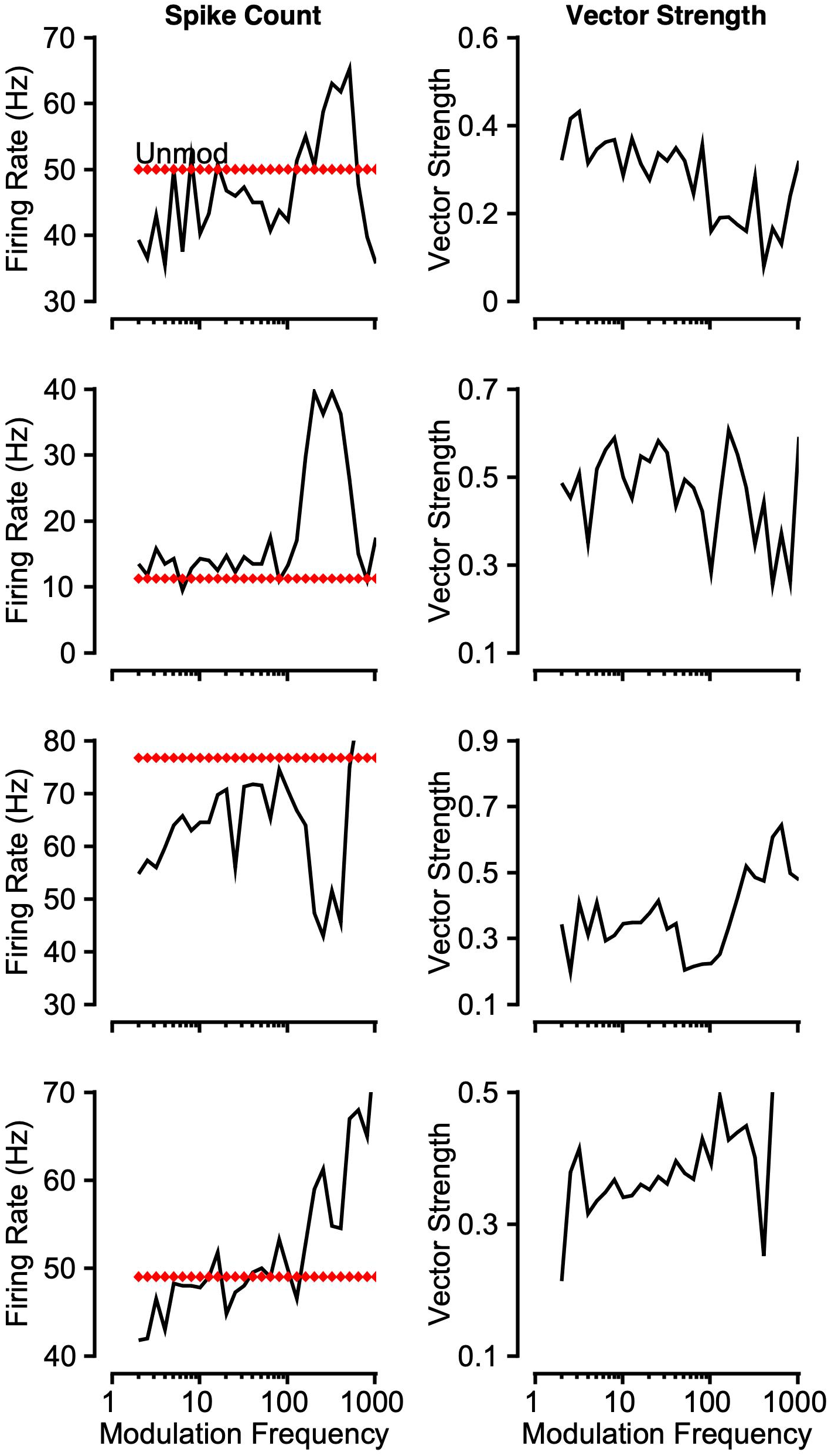
Examples of rate tuning in the CN. Each row shows a rate MTFs (left) and VS-based MTFs of example neurons that exhibited rate-tuning in the VCN. Format is the same as in Figure 4.

### Modulation Transfer Functions in the Inferior Colliculus (IC)

Example rate-based and spike-timing (vector strength) based MTFs from the IC are presented in Figure 6. Most neurons displayed both rate and VS tuning to MF, in contrast to the CN, where less rate-tuning was present. We observed mainly band-enhance/band-pass, high-pass, and band-suppress/band-reject response types, consistent with previous literature (Henry et al., 2016; Nelson and Carney, 2007). The entire tonotopic axis was sampled, as evidenced by the distribution of characteristic frequencies, shown as an inset in the top panel of Figure 6. As is commonly observed, VS tuning to AM was also observed, which facilitated neurometric discrimination analysis described in the following section.

**Figure 6.**
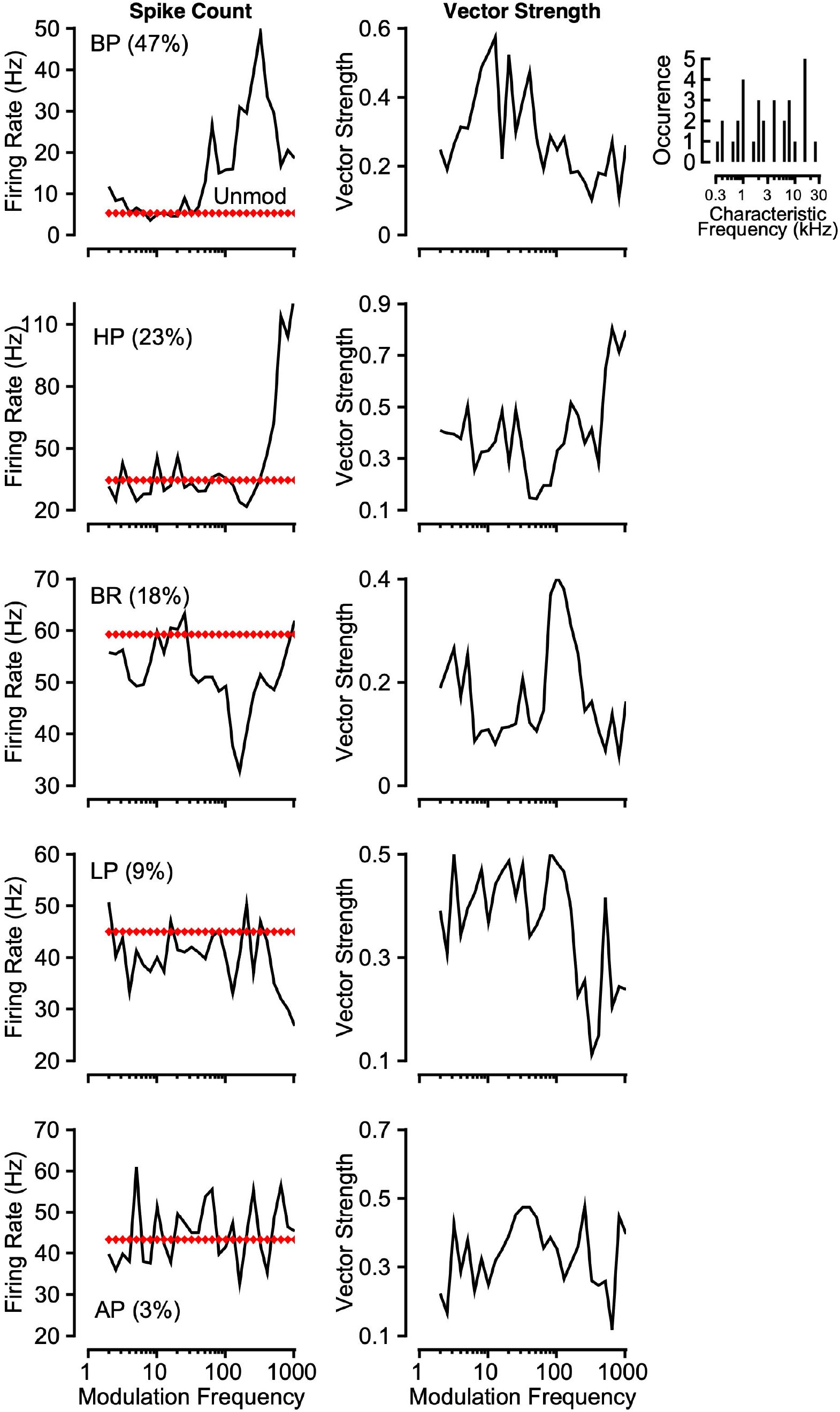
Modulation transfer functions in the inferior colliculus. Each row shows an example neuron’s responses to amplitude modulated noise as a function of the modulation frequency with rate and vector-strength (VS) based MTF on left and right respectively. Neurons exhibited conventional shapes (Band-pass, band-reject, high-pass, low-pass, all-pass). The percentage of each rate-MTF shape encountered is displayed in the figure legend of each panel. Format is the same as in Figure 4.

### Neurometric measures of temporal discrimination based on single-unit activity

Rate and vector strength tuning to AM in the CN and IC permitted the derivation of neurometric measures of temporal discrimination, as previous studies have done as a function of modulation depth (Henry et al., 2016; Johnson et al., 2012; Sayles et al., 2013). Here this type of analysis is extended to modulation frequency discrimination. Figure 7 shows this analysis for an example neuron in the IC. From either the rate or temporal MTF (Figure 7A and B), ROC analysis could be conducted on the distribution of responses (see Methods). The area under the ROC curve is reported here as Neurometric Probability Correct (Figure 7A’). Neurometric threshold criterion is the conventional value of PC = 0.76 (Mackey et al., 2022; Rocchi and Ramachandran, 2018, 2020). Neurometric functions were fit with a Weibull curve (see Methods) to extract threshold. Figure 7A’ and B’ show the corresponding neurometric functions derived via ROC analysis (shown as inset) on the responses in Figure 7A and B. Figure 7C and D show neurometric discrimination thresholds extracted from single neurons near their best modulation frequency. Thresholds were plotted cumulatively, revealing that IC rate-based thresholds (green) are lower than CN (black) rate-based thresholds (Figure 7C; KS test, *p = 0*.*005*). Neurometric discrimination thresholds based on vector strength were also calculated in the same way. Discrimination based on classical vector strength is overestimated when spike counts are very low, which motivated Yin et al. (2011) to introduce a measure called phase-projected vector strength (VS_pp_). We found no significant differences in VS- and VS_pp_-based neurometric discrimination thresholds (two-sample K-S test, *p = 0*.*5*), likely due to the high spike counts typically observed in the subcortical auditory system (compare solid to dotted traces in Figure 7D). Both VS and VS_pp_ based neurometric thresholds were lower in the IC than the CN, indicating greater sensitivity in the IC (Figure 7D), similar to results from rate-based measures (Figure 7C). Neurometric sensitivity as assessed by rate or vector strength did not obviously differ by frequency response map type (data not shown). However, correlations between response types and neurometric sensitivity are better suited for larger sample sizes (e.g. Henry et al., 2016; Nelson and Carney, 2007).

**FIGURE 7.**
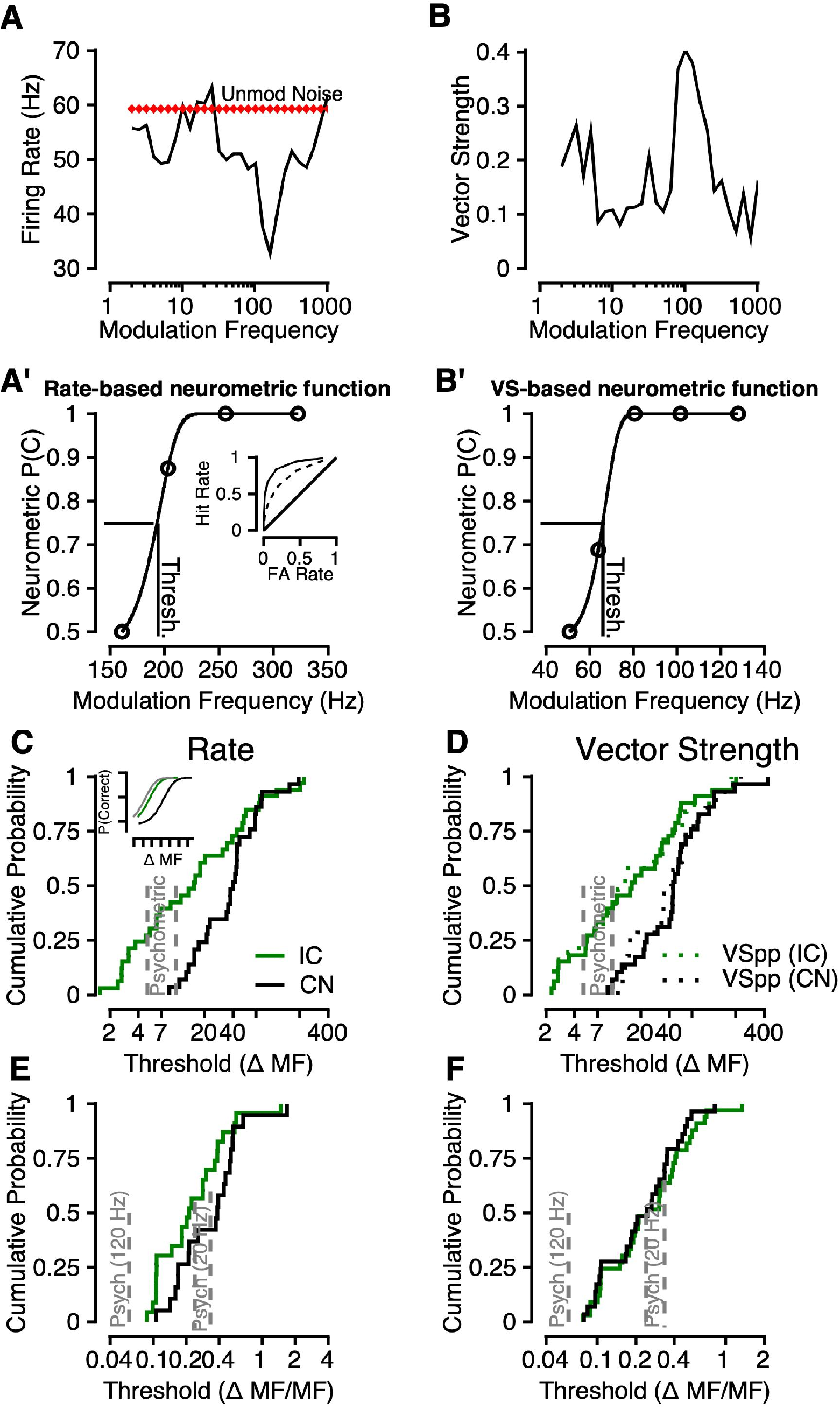
Temporal discrimination based on single-unit activity. **A**. An example band-reject/band-suppress rate-MTF from an IC neuron. **B**. An example VS-MTF from the same IC neuron as in **A. A’**. An example of a derived neurometric function via ROC analysis (see Methods) using the data in A. **B’**. An example of a neurometric function using the data in B. **C**. Firing rate based neurometric thresholds of all units in the sample, plotted cumulatively. IC thresholds are shown in green while CN thresholds are shown in black. **D**. VS and phase-projected VS (VS_pp_) based neurometric thresholds plotted cumulatively in the same format as in C. **E**. Firing rate based neurometric thresholds of all units in the sample, plotted cumulatively and normalized to the standard/comparison modulation frequency used for discriminant analysis. **F**. VS based neurometric thresholds plotted cumulatively in the same format as in E. In all panels psychometric functions are plotted as grey lines.

It was a concern that differences in neurometric sensitivity were being conflated with differences in the range of modulation frequencies to which each neuron was tuned. To address this, neurometric thresholds were normalized to the standard (comparison) stimulus frequency (Figure 7E and F). The IC still displayed lower rate-based thresholds than the CN after normalization, suggesting this effect was not frequency dependent. However, normalized VS-based thresholds in the CN and IC were nearly identical. To relate neurometric sensitivity to psychometric sensitivity, Figure 7 C-F shows monkeys Is and Da psychometric thresholds from the 20 Hz condition in all panels as grey lines. About 40% of IC neurons appear to match or exceed behavioral sensitivity (Figure 9A-B) regardless of which code (rate or VS) is used, while only the most sensitive CN neurons approximate behavioral thresholds. Monkey Is also completed the task in the 120 Hz condition, which is shown in grey in Figure 9 C and Figure 9D. Normalized neurometric thresholds are much higher than monkey Is’ psychometric threshold in the 120 Hz condition (and previous macaque psychometric thresholds at high frequencies). This motivated further analysis of the effect of frequency in the next section.

**FIGURE 8.**
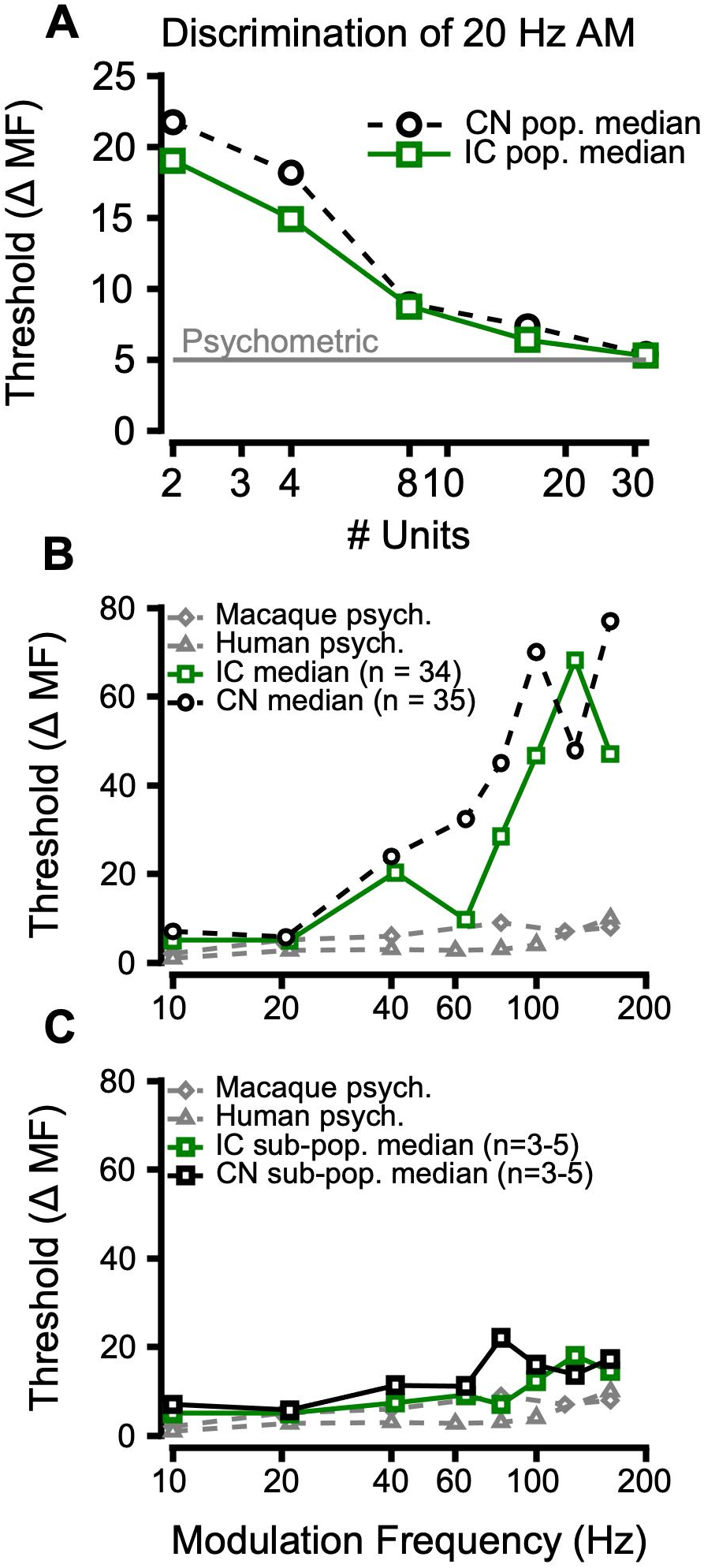
Neurometric thresholds of simulated neuronal populations. **A**. Example trend of neurometric discrimination threshold of indiscriminately sampled populations of neurons (see Methods) as a function of how many neurons were in the population. CN data are shown in black, IC data are shown in green, and behavioral data are shown in grey. **B**. Neurometric discrimination thresholds of indiscriminately sampled neuronal populations in the IC (n = 34 neurons, green trace) and CN (n = 35 neurons, black trace) compared to behavior (grey traces) as a function of the MF of the standard/comparison stimulus. **C**. Neurometric thresholds of small populations (n = 3-5 neurons) compared to behavior. Colors match panel B.

**Figure 9.**
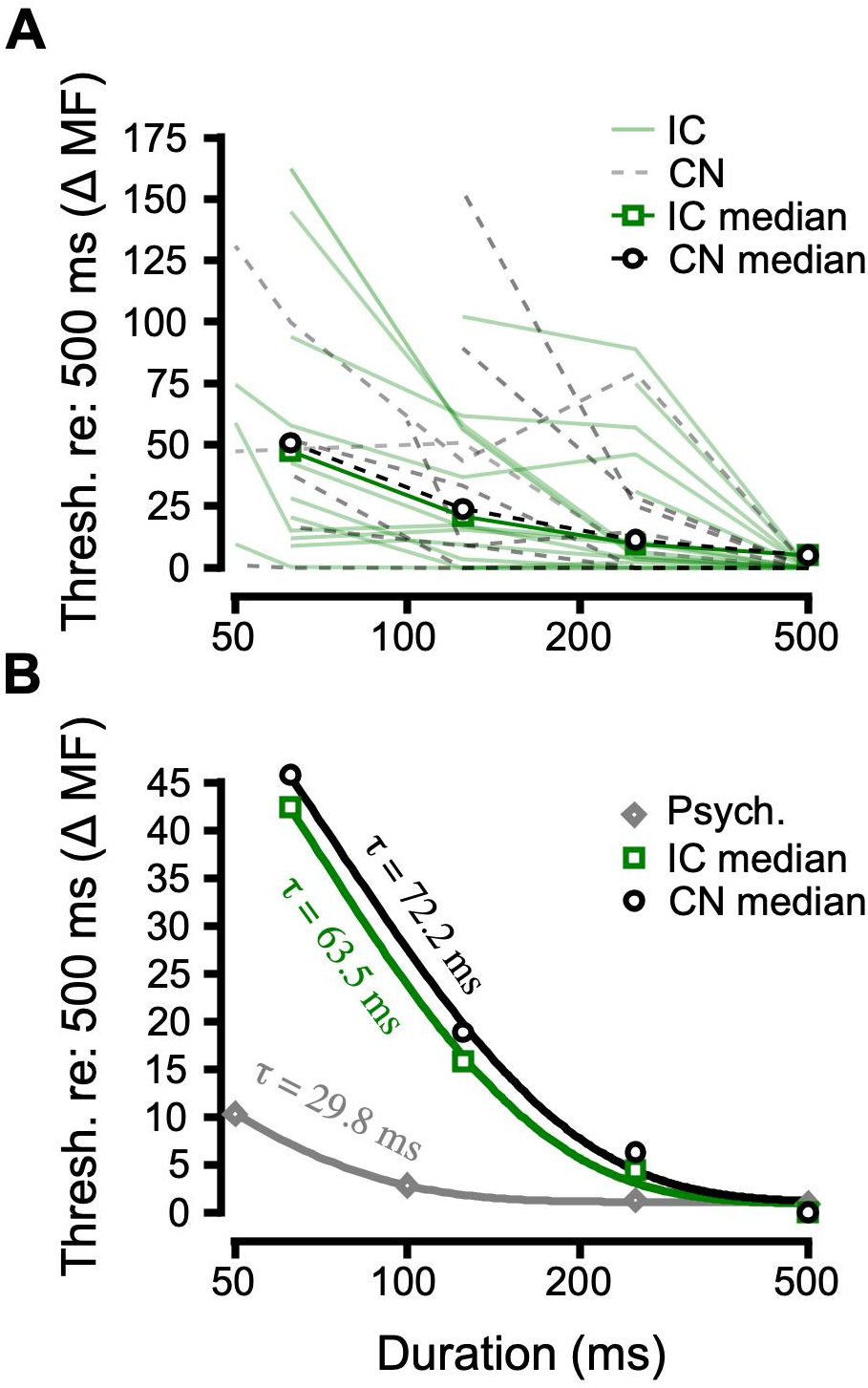
Single neuron thresholds as a function of duration. **A**. All neurometric thresholds plotted as a function of stimulus duration (see Methods) for the IC (green) and CN (black dashes). Thresholds generally increased as duration decreased, as evidenced by the change in the median thresholds (green and black dashes with symbols). **B**. Median IC (green), CN (black), and mean psychometric (grey) thresholds plotted as a function of stimulus duration, and fit with three term exponential functions. The time constant (τ) of each exponential function is listed near each trace.

### Neurometric discrimination based on simulated population activity

Interpretation of single-unit data requires implicit assumptions about how population activity is read out. Previous studies have addressed this by pooling responses in various ways to make explicit how single-unit responses may be combined and related to psychometric measures. Johnson et al. (2012) described an “across-cell method,” by which responses are sampled indiscriminately, with replacement, across the population (see Methods). This permitted the analyses in Figure 8A, which shows that as the number of units in the population increases, the median neurometric discrimination threshold of the population gradually approaches psychometric threshold (Δ5 Hz). While the IC (green squares) was initially more sensitive, CN (black circles) and IC sensitivity converged at 8 units. Once the population size reached 30, both the IC and CN exhibited the same threshold as derived from macaque behavior. This similarity between simulated population and behavioral sensitivity held true for 10-20 Hz, but not for higher frequencies (Figure 8B; macaque behavioral data from Moody (1994) and the present study; human data from Formby (1985)). However, by including only the few neurons tuned to the frequency of interest, simulated populations could exhibit the same sensitivity as behavior (Figure 8C). This necessity of a more complex form of decoding for higher frequencies has been found in a study of gerbil auditory cortex (Penikis, 2020).

### Effects of duration

Neurophysiological studies of envelope perception often use stimuli with a constant duration (Henry et al., 2016; Johnson et al., 2012; Sayles et al., 2013; but see Yao et al., 2020). However, psychophysical studies have gained insight into auditory temporal integration by manipulating duration in AM processing tasks (Dau et al., 1997; Lee, 1994; O’Connor et al., 2011; Sheft and Yost, 1990). This question is of particular importance in the context of macaque studies, as macaques and budgerigars may exhibit unique similarity to humans in temporal integration (Mackey et al., 2021a). This motivated us to evaluate the effects of duration on performance in the AM frequency discrimination task previously discussed and explore its neural correlates. Figure 9 shows discrimination thresholds as a function of duration. As duration became shorter, macaques Is and Da exhibited changes in discrimination threshold consistent with temporal integration. Similarly, neurometric discrimination thresholds changed as the time-window used for spike-count analysis was changed (see Methods). The change in threshold as a function of duration is often quantified by the time constant (τ) of a three term exponential function (Mackey et al., 2021a; O’Connor et al., 1999), and is inversely related to the rate of integration. Thus, Figure 9 shows that CN neurons had, on average, slightly longer time constants than IC neurons, and neuronal time constants were longer than behavior. This suggests that behavioral measures of the temporal integration of AM may more closely resemble changes in higher brain structures (see Discussion). In the absence of data from such structures, some of which may serve as neural integrators of sensory evidence, we modeled the evidence accumulation process using a drift-diffusion model (DDM).

The DDM has been used to formalize assumptions about the evidence accumulation and decision-making process across many different psychological paradigms (de Gee et al., 2020; Liu et al., 2015; Murray et al., 2020; Ratcliff and Murdock, 1976; Ratcliff et al., 2018; Tsunada et al., 2015, 2019), and psychoacoustic studies have modeled AM perception at the computational level (e.g. Dau et al., 1997), but connections have not been made between AM perception and the DDM. In this study, having characterized temporal integration using signal detection theory at the sensory evidence level (the IC and CN) we then sought to computationally characterize neural integration of that sensory evidence using a DDM (see Methods, Figure 10). Drift rate was allowed to vary with noise modulation frequency and duration (Figure 10A and B), which lead to predicted changes in accuracy that resembled empirical psychometric functions (solid lines vs. dashed lines Figure 10 C and D). Predicted discrimination thresholds and empirical discrimination thresholds (Hit Rate = 0.5) were nearly identical (Figure 10E and F). The goodness of fit was quantified with a posterior predictive check (see Methods) confirming that the 95% confidence intervals around the model estimates overlapped with the behavioral data. Monkeys Is and Da exhibited notable individual differences in performance that the model reproduced. The model also accounted for changes in speed (Figure 10G and H) as a function of duration. Combined with the single-unit analysis, these results provide an account of AM perception at the computational and neurophysiological levels and constrain future neurophysiological studies of how acoustic evidence is integrated (see Discussion).

**Figure 10.**
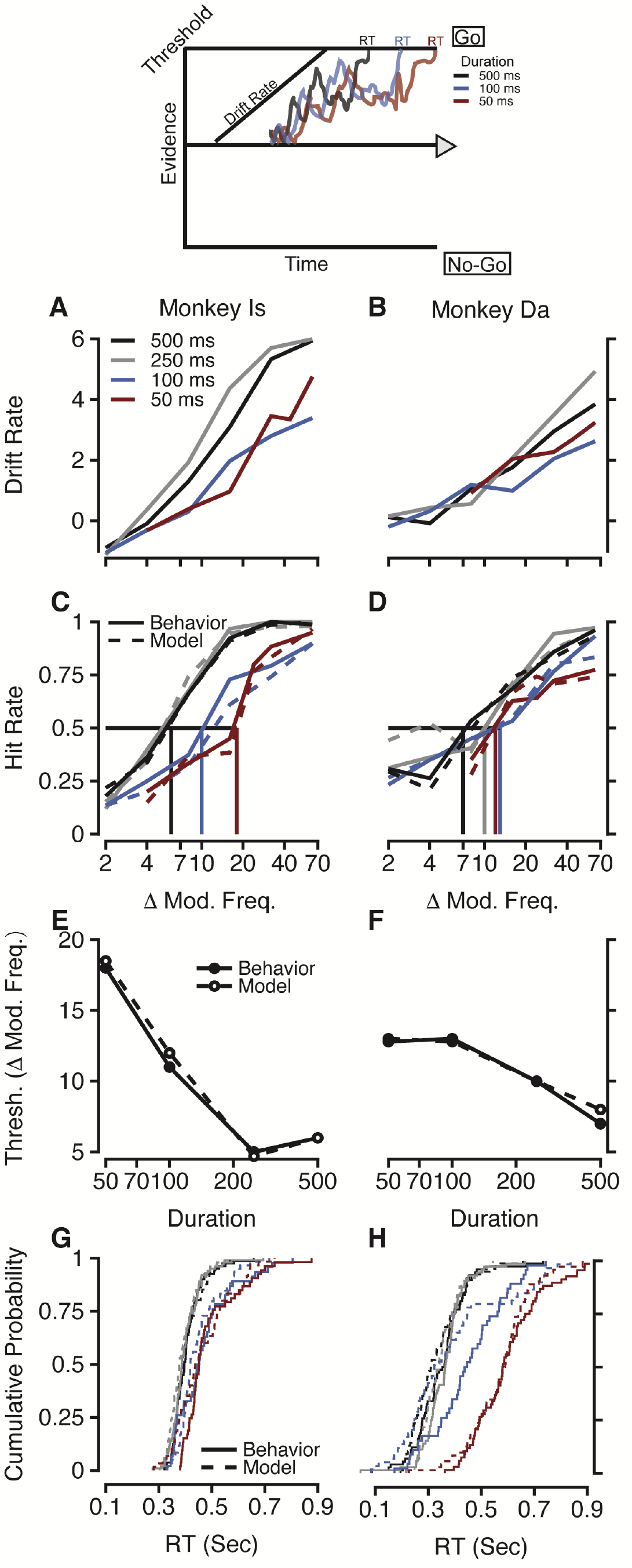
A hierarchical drift-diffusion model (DDM) of temporal discrimination performance. Top, an illustration of how manipulation of drift rate of a diffusion process can lead to predicted changes in reaction time in a Go/No-Go task. **A, B**. Drift rates for stimuli of different durations (shown as different colors), plotted as a function of modulation frequency, for monkeys Is (A) and Da (B). **C, D**. Empirical (solid) and simulated (dashed) psychometric functions produced by the DDM using drift rates shown in panels **A, B. E, F**. Empirical (solid) and simulated (dashed) discrimination thresholds functions produced by the DDM plotted as a function of duration using drift rates from panels **A, B. G, H**. Cumulative reaction time distributions from each monkey (solid) and from the DDM (dashed), with different durations represented by different colors, shown in panel A.

## Discussion

The present results confirm that there is an increased prevalence of rate-coding of AM as the primate auditory neuraxis is ascended, which may support sound envelope perception. However, a few of the results presented here may indicate need for reconsideration of the existing literature on how the primate auditory system processes AM. This partially stems from the fact that, to the authors’ knowledge, there is only a single study on how the neurons in the primate CN encode AM, and no such studies of the primate IC. The CN study used an anesthetized preparation, and found that there was no rate encoding of AM (e.g. all-pass rate MTFs) in the cochlear nucleus (CN) of multiple primate species (Rhode et al., 2010). This differs from the present results, where rate-tuning was encountered despite the small sample size. The simplest explanation for this difference seems to be the use of anesthesia, which is known to impact sound encoding (Noda and Takahashi, 2015; Ramachandran et al., 1999; Schumacher et al., 2011). However, rate-tuned CN neurons have been found in other species, even under anesthesia (Joris et al., 2004). These rate-tuned neurons generally display onset-chopper PSTHs. However, in our admittedly small sample, we did not encounter onset-choppers, despite encountering rate-tuning. It seems unlikely that this is sampling bias, as few onset-choppers were found in a larger sample (92 neurons) from this lab (Mackey et al., 2022; Ramachandran, 2018). This difference could be due to the use of an awake experimental preparation, the fact that the macaques had experience in AM processing tasks, or a unique aspect of primate CN neurophysiology. The last conclusion is supported by anatomical studies of primate CN anatomy (Adams, 1986; Kavanagh Moore, 1980b; Moore et al., 1996; but see Rubio et al., 2008).

The CN results presented here also differ with respect to how they relate to perception. The finding that average cochlear nucleus (CN) neuronal spike-timing is not as sensitive as behavior (see Results), contrasts a previous study in anesthetized chinchilla CN--the only study where comparable neurometric analyses were done using CN data (Sayles et al., 2013). Sayles et al. found that CN spike-timing, on average, was as sensitive as chinchilla behavior, and they thoroughly addressed necessary methodological caveats in their discussion (e.g., anesthesia). It’s possible that the use of anesthesia is responsible for the differences between their data and the present data. This difference could also represent a species difference, as chinchillas and primates display perceptual differences in temporal discrimination (See Figure 3; Moody, 1994).

Many studies have addressed questions about what neural code present in the IC supports envelope perception (Henry et al., 2016, 2017; Lorenzi et al., 1995; Nelson and Carney, 2004, 2007). Two studies that are very comparable to the present results have suggested rate responses, even optimally pooled across a large sample, are insufficient to explain AM detection and depth discrimination performance (Henry et al., 2016; Nelson and Carney, 2007). Given this, it is surprising to consider that both spike rate and timing measures in the IC provided a close match to macaque and human AM discrimination performance in the present analysis. Similar hierarchies in temporal encoding have been found in a study of IC, medial geniculate body, and primary auditory cortex (Asokan et al., 2021), which characterized how progressively longer timescales are encoded in these regions. These results (Figure 8) are consistent with such a hierarchy, and build on our previous finding that IC responses provide more reliable estimates of psychometric threshold and slope, compared to CN responses, and that IC neurons exhibit significant choice-probabilities (Mackey et al., 2022). This neurometric-psychometric correlation supports the notion that the greater prevalence of band-enhanced and band-suppressed rate profiles in the midbrain may have significance for perception that was previously unclear due to the greater sensitivity of spike-timing found in Henry et al. (2016) and Nelson and Carney (2007). This conclusion is consistent with previous IC model comparisons to human AM detection (Lorenzi et al., 1995).

The results presented here are also interesting in the context of previous AM studies in macaques, which have exclusively focused on the cortex. A previous study in A1 found that single neuron rate or timing, on average, were less sensitive than macaque AM detection behavior; consistent with the lower-envelope principle (Johnson et al., 2012). This is surprising given that the present results suggest nearly half the single neuron sample recorded in the IC matched behavior in rate and vector strength. The finding that average single neuron sensitivity could not explain behavior, led Johnson et al. to pool responses in the same way as the present study. Their analysis suggested 25 neurons’ rate responses were sufficient to account for behavior. This was the case particularly for low modulation frequencies (10-20 Hz) where vector strength (single neurons or pooled responses) was insufficient to explain AM detection. Sensory evidence necessary for the *discrimination* (as opposed to detection) of such low (“flutter”) frequencies has been suggested to reside in A1 in the form of a rate code (based on macaque data: Lemus et al., 2009b). However, the results presented here suggest a substantial degree of perceptual AM sensitivity could be inherited from subcortical stations in the form of a rate or spike-timing code.

These differences from previous CN and IC studies must be taken with a caveat: the onset response has been excluded in many previous studies (Beitel et al., 2003; Henry et al., 2016; Johnson et al., 2012; Sayles et al., 2013), though, interestingly, Johnson et al. (2012) report that “including the onset response did not substantially alter the results.” Some more recent studies of primate auditory cortex have included the onset response, which facilitates comparison to the current results. These studies also used a modulation frequency decoding framework, further aiding comparison to the present results (Downer et al., 2021; Hoglen et al., 2018). Hoglen et al. (2018) report that decoding of modulation frequency exhibited similar patterns across 0-750 ms and 250-1000 ms time-windows, which may indicate that the onset response does not alter results in the present report, which is consistent with the remarks in Johnson et al. (2012). In this report, the onset response was included to facilitate comparison with these previous publications and with the behavioral experiments, where presumably the macaques may use the onset response. This also enabled us to make comparisons to experiments where the duration of the stimulus was reduced to 50 ms, where presumably only some form of onset response is available. Psychometric thresholds at short durations were substantially lower than most neurometric thresholds, suggesting that macaques can use information present in the onset response. The present results are not the first to suggest this, as a previous study provides strong evidence that the onset response in the IC is of great utility for localization performance in reverberant environments (Devore et al., 2009).

The effects of duration in this study extend work on auditory temporal integration to an AM discrimination paradigm, as opposed to more common tone detection paradigms (Costalupes, 1983; Heil et al., 2017; Mackey et al., 2021a; O’Connor et al., 1999; Plomp and Bouman, 1959). Temporal integration of AM stimuli has been explored almost exclusively at the psychophysical level, and mostly in humans (Dau et al., 1997; Lee, 1994; O’Connor et al., 2011; Sheft and Yost, 1990), though one study has reported causal evidence that parietal cortex is involved (Yao et al., 2020). Yao et al. (2020) used a gerbil model, which permits the use of powerful genetic techniques. However, our previous work found that macaques and budgerigars exhibit unique similarity to humans in auditory temporal integration (Mackey et al., 2021a), which highlights the value of exploring these questions in a variety of animal models. The behavioral time constant reported here (30 ms) is almost identical to our previous report (Mackey et al., 2021a), and is much lower than the IC and CN (Figure 9), which suggests IC and CN integrate AM more slowly than macaques. A related question is how downstream regions of the brain integrate sensory evidence. Posterior parietal cortex (PPC) is one candidate integrator of acoustic evidence (Yao et al., 2020), and sensory evidence more generally (e.g. Cohen et al., 2004; Huk and Shadlen, 2005). Alternatively, the superior colliculus (SC) could serve this role, as multiple studies of spatial processing have found (Jay and Sparks, 1987; Rajala et al., 2017; Wallace et al., 1996), and there is causal evidence for its role in evidence accumulation in the visual domain (Jun et al., 2021). In the absence of neural data from one of these regions in this study, a DDM was used to characterize the integration process at the computational level, providing constraints on future neurophysiological studies in regions like PPC or SC. Specifically, the model results suggest that the effects of modulation frequency on the rate of integration decrease as duration decreases. This is indicated by the slope of the drift rate functions in Figures 10A and B, which increase as duration decreases. Candidate neural sites of integration should display this key aspect of auditory temporal integration.

## Acknowledgements

The authors would like to acknowledge Bruce and Roger Williams for fabrication of hardware, Mary Feurtado for assistance with surgical procedures, and Dr. Jane Burton collecting neurophysiological data. The study and RR were supported by research grant NIH R01 DC 015988 (MPIs R. Ramachandran and B. Shinn-Cunningham) and NIH RO1 DC 11092 to RR. SH was supported by NIH T35 DC 008763 (PI Linda J. Hood). CM was supported by the Ruth Kirchstein pre-doctoral fellowship from NIDCD (F31 DC 019823-01).

